# Rising above the noise: The influence of population dynamics on the evolution of acoustic signaling

**DOI:** 10.1101/2022.09.02.506422

**Authors:** Megha Suswaram, Uttam Bhat, Justin D. Yeakel

## Abstract

Acoustic signaling is employed by many sexually reproducing species to select for mates and enhance fitness. However, signaling in dense populations can create an auditory background, or chorus, which can interfere with a signal receiver’s phonotactic selectivity, or the ability to distinguish signals. Feedback between the strength of an individual’s signal, phonotactic selectivity, and population size, may interactin complex ways to impact the evolution of the signaling trait within a population, potentially leading to the emergence of silence. Here we formulate a general model that captures the dynamic feedback between individual acoustic signalers, phonotactic selectivity, and the population-level chorus to explore the eco-evolutionary dynamics of an acoustic trait. We find that population dynamics has a significant influence on the evolutionary dynamics of the signaling trait, and that very sharp transitions separate conspicuous from silent populations. Our framework also reveals that increased phonotactic selectivity promotes the stability of signaling populations. We suggest that understanding the relationship between factors influencing population size such as environmental productivity, as well as factors influencing phonotactic selectivity such as anthropogenic noise, are central to understanding the complex mosaic of acoustically signaling and silent populations.

## 1 INTRODUCTION

Acoustic signaling is the primary mode of communication shared by roughly 8.7 million species ranging from arthropods to mammals (Chen and Wiens, 2020), inhabiting both terrestrial and marine environments. While acoustic signaling serves many functions, one of central importance is to attract potential mates. Several characteristics of acoustic signalling, such as the length of the signal, repetition rate, frequency, amplitude, pitch, and decibel level can be used to create a meaningful and distinct auditory signal (Desutter-Grandcolas and Robillard, 2011; Rowe, 1999; Nityananda and Balakrishnan, 2021). While acoustic signaling is efficient, it does not come without costs. For instance, conspicuous signalers, or individuals producing acoustic signals far above the population mean, while easily located by potential mates (de Crespigny and Hosken, 2007) can also be located by potential predators (Sakaluk, 1990; Endler, 1992) and parasites (Zuk and Kolluru, 1998). Moreover, conspicuous signaling can be energetically taxing, thus depleting energetic reserves that may otherwise be invested in growing and maintaining somatic tissues, or directly invested into offspring. Furthermore, the time spent signaling takes away from that spent foraging (Abrahams, 1993a). Finding the balance between the reproductive rewards from signaling, while both avoiding predators and maintaining adequate energetic reserves (Abrahams, 1993b; Ellers, 1996), is a central challenge for organisms specializing in this mode of communication.

Signal-producing traits are subject to selection, and depending on the costs and benefits may intensify or diminish signaling over evolutionary time. In some cases, changes in the trade-offs introduced by acoustic signals can lead to both rapid evolution as well as disruptive selection, driving the inherent acoustic diversity within particular phylogenetic groups such as Pacific field crickets (Tinghitella et al., 2018). Within a population, the signaler interacts not only with potential signal receivers, but also with its acoustic environment (Balakrishnan et al., 2014; Wells and Schwartz, 2007; Gerhardt, 2001; Nandi and Balakrishnan, 2016). By producing a signal, the signaler in turn modifies the acoustic environment, which feeds back to affect the costs and benefits associated with signaling among conspecifics. In this sense, acoustic signaling can be thought of as an immediate form of niche construction (Balakrishnan et al., 2014; Schwartz and Bee, 2013), as signaling behaviors directly alter the acoustic environment. As competing signalers must engage this changing environment directly in order to rise above the noise, this feedback may result in further alteration to the acoustic environment. This will alter the fitness benefits of the signaling, changing the shape of the fitness landscape for all individuals in the population (Hartbauer et al., 2014).

The magnitude of the acoustic background produced by local signalers, here and henceforth referred to as the chorus, is influenced by both the traits of individual signalers as well as the number of signalers within the population. If individual signalers are conspicuous, and there are many of them, then the chorus is conspicuous. At the extreme, if most individuals in the population are silent, so is the chorus. In addition to the influence of individual signallers on the chorus, the size of the population giving rise to the chorus is expected to directly influence the potential reproductive advantage attributed to a small change in an individual’s signal. If the population is large, a minor increase in an individual’s signal is expected to have negligible effects on the individual’s reproductive gain. If the population is small, a minor increase in an individual’s signal may carry with it larger reproductive advantages, as it is easier for signal receivers to target and reward the signaler when there are fewer signalers to navigate (Gerhardt and Klump, 1988; Niemelä et al., 2019). The ability of a receiver to discern between individual signalers, known as phonotactic selectivity, increases the sensitivity of reproductive gain to increases in signal (Gerhardt, 2008; Jang et al., 2010). That phonotactic selectivity is density-dependent means that the eco-evolutionary dynamics of acoustically signaling populations are expected to interact across similar timescales. The potential dynamic outcomes of such a system may thus have consequences with regard to whether signaling traits are reinforced over evolutionary time, or whether they are lost (Safran et al., 2013).

Here we formulate a general model that captures the dynamic feedback between individual acoustic signalers, the population-level chorus, and the eco-evolutionary consequences with respect to a single continuous acoustic trait. Our framework provides insight into a) the complex interaction between population density and phonotactic selectivity in determining the fitness landscapes of signalling populations b) the influence of reproductive rewards on the evolutionary transitions between conspicuous and silent populations, and c) the effects of increasing energetic costs on the evolution signaling. Together, we demonstrate how the inherent feedbacks between the reproductive advantages associated with signaling against changes in population density and strength of the chorus can position some populations to evolve towards mate-signaling and others towards silence. As the benefits of signaling vary across heterogeneous environments, these feedbacks may contribute directly to the complex geographic mosaic of acoustic strategies observed among closely related taxa.

### 1.1 Model Framework

We consider a population of organisms using acoustic signaling to attract potential mates. Individual fitness is thus a trade off between the reproductive advantages of mate-signaling, and the acquisition and maintenance of energetic stores to ensure survival. Here and henceforth, we will use field crickets as our exemplary system, however we emphasize that our framework is general and relevant to any population that must balance the energetic costs of signaling against a background of signaling conspecifics. We assume that individual fitness is a function of a single quantitative character, which we specify here as the syllable rate *z*, scaled to range between *z* = 0 (silence) to *z* = 100 (maximal signaling). We note that this trait could easily map onto any aspect of an acoustic signal with the only requirement being that it has a temporal and/or energetic cost. The potential reproductive gain of an individual’s signal is determined by how well it can be distinguished from the chorus, which in this case is represented by the mean syllable rate of the population 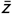, which itself evolves over time. We thus track the eco-evolutionary dynamics of the population by evaluating the fitness of individuals with respect to an evolving chorus of signalers, where energetic investment in a higher syllable rate relative to the chorus is met with an opposing divestment in those behaviors needed to acquire and maintain the organism’s energetic stores and avoid a reduction in fitness. We capture the population dynamics and evolution of *z* with a discrete-time individual-based model, where we assume non-overlapping generations.

### 1.2 Individual fitness of a signaler

In a sexually-signaling population, reproductive gain is increased when an individual broadcasts a signal that can be distinguished by potential mates from the sensory background. Among crickets, a population of individuals signaling together with variable syllable rates *z* forms the chorus 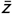. In an environment devoid of external acoustic interference, this mean syllable rate sets the reproductive standard for all signaling individuals within the population: individuals with 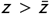 are assumed to have a reproductive advantage over those with 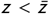. We assume that reproductive gain is described by a sigmoidal function, where *r*_min_ represents the non-zero but minimal reproductive reward obtained by silent individuals, increasing to the maximum reproductive reward *r*_max_ obtained by individuals with syllable rates far above the chorus 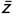. The chorus 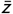 describes the location of the step transition of the reproductive gain function, which changes dynamically over time as the population evolves (Fig. 1).

**FIGURE 1.**
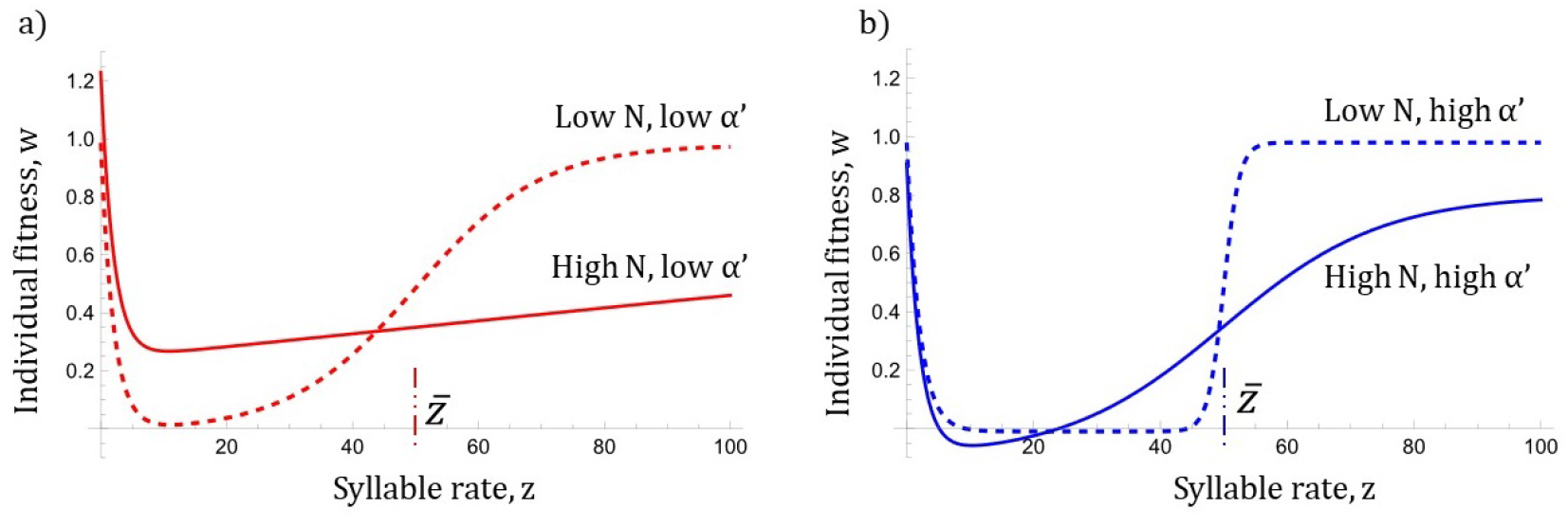
Individual fitness 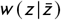 as a function of low (dashed line) and high (solid line) population densities *N*, and with respect to a) low (red) and b) high (blue) auditory sensitivities. If auditory sensitivity is low, phonotactic selectivity remains limited even at low population sizes. If auditory sensitivity is high, phonotactic selectivity is enhanced at low population sizes, resulting in steep reproductive gains about the chorus 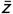.

In sparsely populated habitats where individual calls are more easily distinguished, even a small change in an individual’s syllable rate *z* could result in a significant reproductive advantage if it is even slightly above the chorus, which in this case would be the product of a small number of conspecifics. This corresponds to 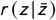 with a steeper slope from *r*_min_ to *r*_max_, where the half-saturation is located at the chorus mean 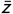 (Fig. 1b). In contrast, in densely populated habitats, a small change in *z* near or above the chorus would be expected to have a much smaller effect on individual fitness. This corresponds to 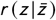 with a shallower slope from *r*_min_ to *r*_max_ about the chorus mean (Fig. 1). For example, mating success among male signalers tends to vary inversely with population size among crickets (Niemelä et al., 2019), suggesting that a given signal provides greater reproductive gain when there are fewer competitors to cloud the field. The steepness of 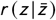 about 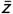, given by *α*(*N*), describes the phonotactic selectivity of the population. In this context, phonotactic selectivity provides a qualitative measure of the auditory discerning ability of the receivers between signalers, where a high value (steep) means that receivers can easily discern between signaling individuals, whereas a low value (shallow) means that receivers cannot easily discern among signaling individuals. As phonotactic selectivity changes inversely with population density, we assume that *α*(*N*) = *α*′/*N*, where *α′* describes the sensitivity of phonotactic selectivity to changes in the population.

As *α′* increases, the effects of phonotactic selectivity become exaggerated, such that smaller population sizes elicit sharper increases in reproductive gains with small increases in *z* above the chorus 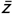 (Fig. 1). This auditory sensitivity *α′* is thus a product of the anatomical signal-receiving equipment carried by the signal-receivers, and is known to be physiologically linked to changes in neural responsiveness (Kostarakos and Hedwig, 2012). Sensitivity to auditory signals is expected to vary from species to species across different environments. For example, among two species of North American gray tree-frogs (*Hyla chrysoscelis and H. versicolor*), females of *H. versicolor* both possess auditory receiving and processing systems that are more sensitive to external stimuli and demonstrate greater selectivity in mate choice (Gerhardt, 2008), in this context implying a higher value of *α*′. Throughout, we examine how increases in auditory sensitivity, promoting greater phonotactic selectivity at low population densities, alters the fitness tradeoffs associated with signaling, and to what extent this might impact potential eco-evolutionary outcomes. Together, the reproductive component of individual fitness is described as

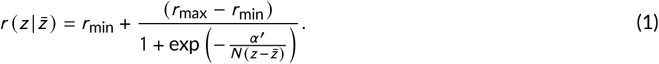

Acoustic signaling entails significant energetic and temporal demands (Hoback and E. WAGNER JR, 1997; Prestwich, 1994). As more energy and time is invested into signaling, the individual has less to invest into foraging, somatic maintenance, and avoiding predation, all of which increases the risk of mortality (Burk, 1988; Casagrande et al., 2016). As such, we assume that silent individuals are subject to a lower per-capita mortality rate, *δ*_min_, whereas maximally signalling individual experiences a higher per-capita mortality rate, *δ*_max_. Mortality as a function of *z* is then written as

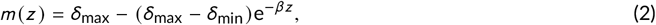

where *β* determines the steepness with which costs increase with higher *z*. Changes in *β* from population to population may capture differences in environmental productivity, where increasing energetic investment in environments with scarce resources may result in larger increases in *m*(*z*) at high signal rates. In natural systems, cricket mortality is highly environmentally dependent (Gutiérrez et al., 2020; Centeno Filho et al., 2021), where in arid regions with limited resource availability, small changes in syllable rate can drastically increase mortality (Gray and Eckhardt, 2001; Dougherty, 2021). When resources are more abundant, individuals have greater energetic latitude and individuals may be more likely to to adopt costly signals (Wagner Jr and Hoback, 1999; Scheuber et al., 2003).

The fitness of an individual signaling with rate *z* is computed by its reproductive fitness, modulated by carrying capacity, *K*, minus the fitness costs of mortality, together given by

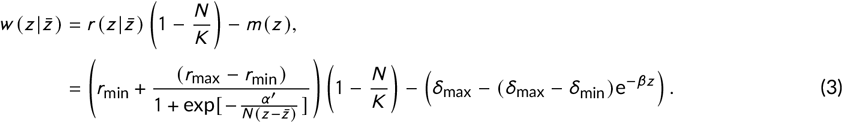

The shape of the fitness function reveals a peak and a plateau, separated by a fitness valley as a function of *z* (Fig. 1). The peak emerges at *z* = 0, or silence, while the variable plateau emerges at high syllable rates, or *z* ≫ 0. While the fitness peak at *z* = 0 reflects that of silent individuals, the fitness plateau at high *z* represents the reproductive gains associated with conspicuous signaling, the value of which results from the trade-off between the energetic costs and reproductive rewards of signaling. As *z* evolves dynamically over time, the shape of the fitness trough is also dynamic, which has particular significance for the evolution of acoustic signaling within the population. Phonotactic selectivity, shown as *α*(*N*) = *α′*/*N*, changes with population size to determine the size and steepness of the fitness trough (Fig. 1). We consider model results where auditory sensitivity *α′* is set to both both low and high values, corresponding to species that are ill- and well-equipped to distinguish individuals apart from the chorus. For simplicity we classify individuals and the population chorus mean as as silent and conspicuous at 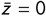 and 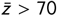, respectively.

### 1.3 Simulation of eco-evolutionary population dynamics

Because an analytical solution for the average fitness of the population 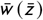 is intractable, we track changes in population abundance and the trait distributions of a cricket population with an individual based model. As such, we numerically track the evolution of the full trait distribution of *z*, denoted by *f*(*z*), over time, in addition to the population size *N* (*t*). Each time-step represents a generation where all adults are assumed to die at the end of each time-step such that generations are non-overlapping. This is generally the case for cricket populations (Masson et al., 2020), however this assumption would not hold for many other acoustically signaling organisms including most bird species. Offspring inherit trait values from parents with variability *σ* such that

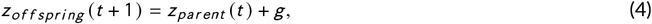

where *g* ~ Norm(0, *σ*), and we set *σ* = 1. The number offspring for each individual *i* with trait *z_i_* is determined by its fitness with respect to the chorus mean, 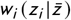, where the sum across reproducing individuals determines the future population size *N*(*t* + 1), such that

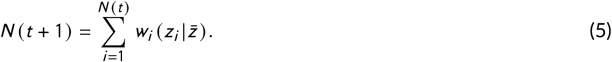

See an illustration of the core framework in Fig. 2. In all instances, the initial population size is set to *N* (*t* = 1) = 1000 individuals and the initial trait distribution is set to *f* (*z, t* = 1) ~ Norm (50,1). The model was simulated for 1000 generations, where we visually confirmed 1000 generations to be more than adequate for calculating steady state conditions.

**FIGURE 2.**
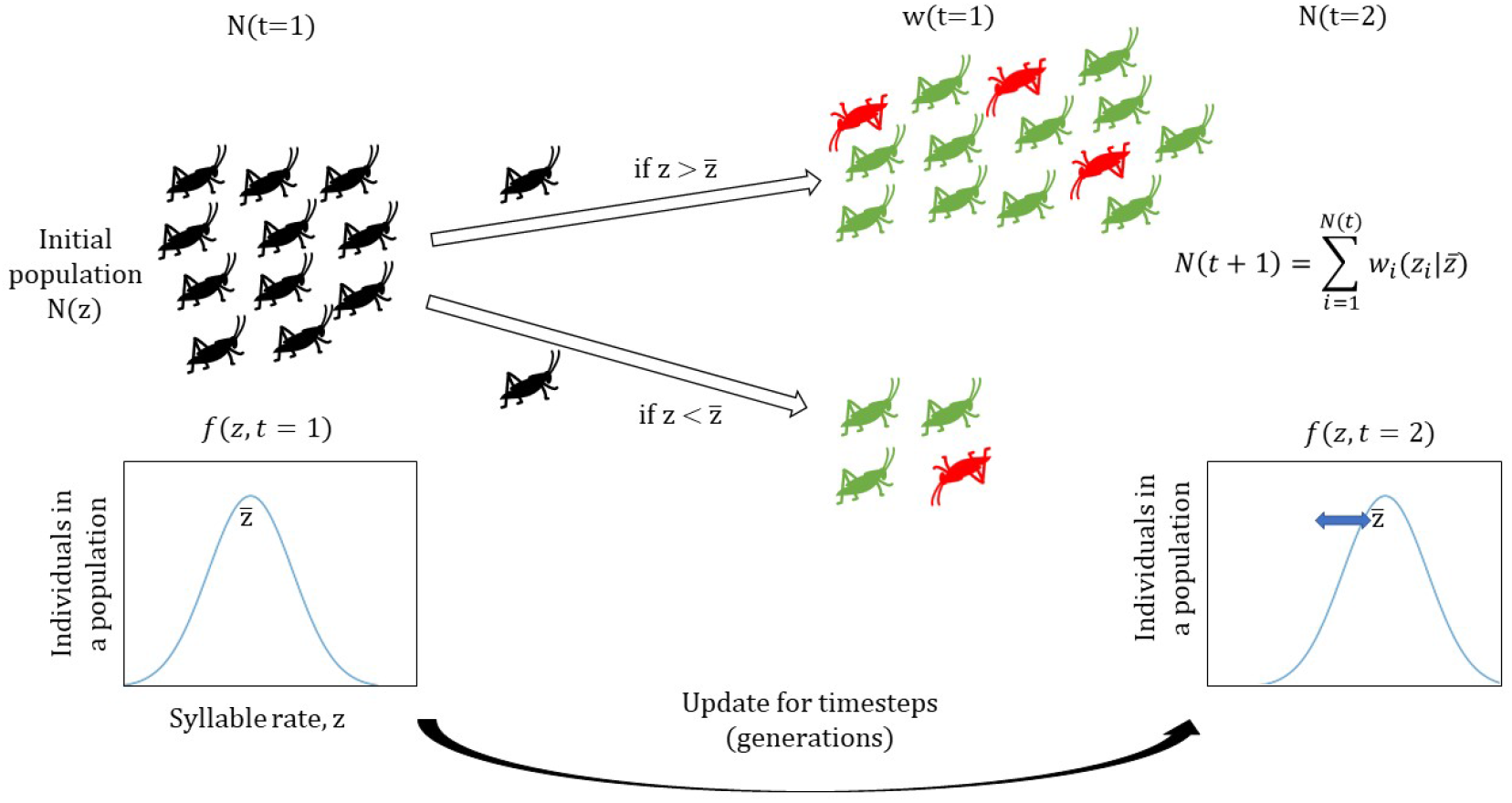
A visual representation of the model framework. N(t=1) is the initial population distribution, where green represents reproduction and red represents mortality. The fitness of the initial population is denoted by w(t=1). N(t=2) represents the population after one time step in which the offspring have slight phenotypic deviation from that of their parents and the chorus mean of the population has shifted.

**TABLE 1.** Parameters used in the simulation

| Parameter | Description | Value/Range |
| --- | --- | --- |
| $z$ | Syllable rate | 0 - 100 |
| $\bar{z}$ | Chorus mean of the population | Var |
| $N$ | No. of individuals of the population | Var |
| $K$ | Carrying capacity | $10^5$ |
| $r_{\min}$ | Minimum reproductive reward without signaling | 1-10 |
| $r_{\max}$ | Maximum reproductive reward with signaling | 1-10 |
| $\alpha(N)$ | Phonotactic selectivity | Var |
| $\alpha'(N)$ | Auditory sensitivity | 1-1000 |
| $\delta_{\min}$ | Minimum per-capita intrinsic mortality | 0.01 |
| $\delta_{\max}$ | Max per-capita mortality rate | 1 |
| $\beta$ | Environmental productivity | 0.5 |

## 2 RESULTS & DISCUSSION

### 2.1 Evolutionary transitions between conspicuous and silence

Our framework reveals the dynamic emergence of both silent and conspicuous signaling regimes, with sharp transitions separating these divergent evolutionary outcomes. Of central importance is the reproductive incentive associated with signaling, which we calculate as Δ*r* = *r*_max_ – *r*_min_. As such, if *r*_max_ is very large with respect to a particular *r*_min_, the potential reproductive gain if the signaler can be distinguished from the chorus is likewise large. If *r*_max_ is only slightly larger than *r*_min_, the potential reproductive incentive is slight. We simulated both the population size *N*(*t*) and trait distributions *f*(*z*) across a range of values for *r*_min_ and *r*_max_ to calculate both chorus mean and population steady states, denoted by 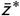 and *N*\*, respectively.

Results of our simulation reveal that silence is the dominant outcome, particularly when the *r*_min_ is relatively high (Fig. 3a). Conspicuous signaling emerges as an evolutionary outcome only when *r*_min_ is very low, and for an intermediate range of *r*_max_. That silence is the dominant outcome when the reproductive gain associated with silence is large is straightforward: the advantages of signaling do not outweigh the energetic costs when silent individuals receive above-minimal reproductive rewards.

**FIGURE 3.**
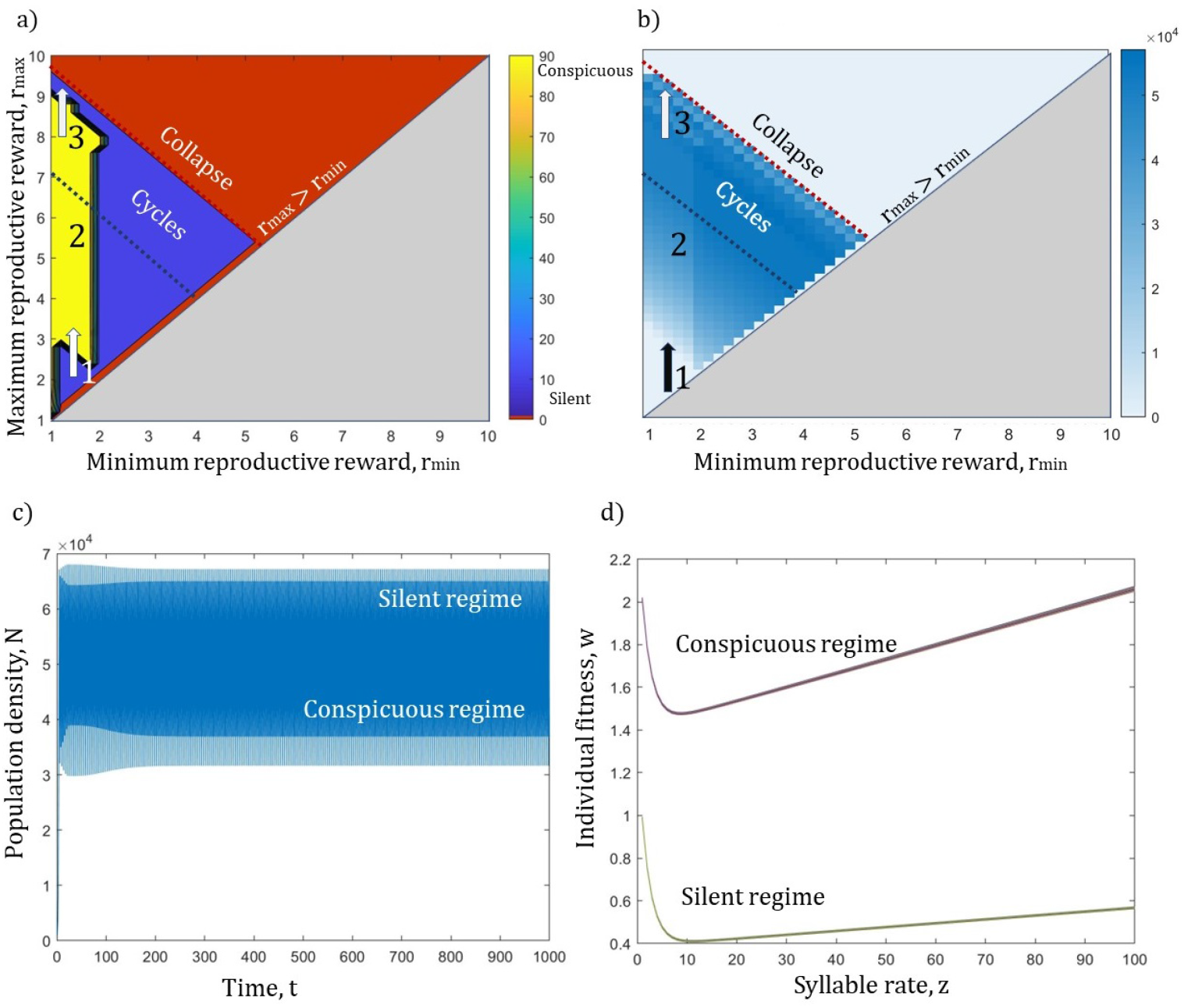
Reproductive incentives drive the evolution of silence or signaling. a) The chorus mean steady states 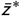 and b) population size steady states *N*\* as a function of minimum *r*_min_ and maximum *r*_max_ reproductive rewards. c) Population steady states in the cyclic regime across 1000 generations, given *r*_max_ = 7 and *r*_min_ = 2. d) Individual fitness 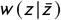 as a function of *z* associated with the population maxima and minima in the cyclic regime as in c). Throughout the auditory sensitivity is assumed to be high, such that *α′* = 800.

To understand the emergence of signaling, one must also take into account the influence of population size, which directly influences phonotactic selectivity, or the ability of a signal receiver to distinguish the signaler from the acoustic background. When both *r*_min_ and *r*_max_ are very low, there are no benefits to signal as Δ*r* is small. Maintaining low *r*_min_, and with increasing *r*_max_ a dynamic threshold is crossed, above which emerges the evolution of conspicuous signaling (transition 1, Fig. 3a,b) At this transition, the small populations densities emerging from these low reproductive rewards enhances the individual fitness associated with conspicuous signaling, as observed in Fig. 1. This gradually increases the chorus, which signalers must surpass to gain the rewards associated with signaling, such that conspicuous signaling is the evolutionary outcome. As *r*_max_ continues to rise, silence re-emerges (transition 3, Fig. 3a,b). Here the reproductive rewards associated with signaling are high resulting in a large number of offspring awarded to signalers. This increased population growth results in an instability, where population fluctuations emerge and give rise to first period-doubling and then the onset of chaos is Δ*r* continues to expand.

That signaling can emerge or be extinguished abruptly among populations is a phenomenon that has been repeatedly observed in natural systems. For example, when signaling incurs additional fitness costs, rapid evolution of silence can result. On the island of Kauai, the presence of parasitoid flies (*Ormia ochracea*) that target and attack signaling oceanic field crickets (*Teleogryllus oceanicus*) resulted in the dominance of a silent flatwing morph within 20 generations (Zuk et al., 2006). For other acoustic groups such as frogs, cicadas and humpback whales, a change in reproductive incentive due to acoustic interference has resulted in the rapid evolution of silence as well as the development of alternative multi-modal communication (Partan, 2017).

The onset of population cycles and eventually chaos when *r*_max_ is very high is a natural result of the logistic relationship assumed for the individual fitness function (Eq. 3), however its effect on the evolution of signaling versus silence is instructive. With the emergence of cycles, the fitness landscape changes abruptly with sharp inter-generational transitions between high and low population sizes (Fig. 3c,d). When populations are large, phonotactic selectivity is weak, such that signalers cannot be easily distinguished from the population’s acoustic background, increasing the relative fitness associated with silence (the silent regime). When populations are small, phonotactic selectivity is much stronger because there are fewer individuals to discern, increasing the relative fitness associated with conspicuous signaling (the loud regime). This quickly fluctuating fitness landscape means that the strength of selection is very weak, such that the across-generational fitness differences are too large for selection to produce an evolutionary response. When population cycles first emerge, this weak selection results in a conspicuous signaling population. Across the period-doubling cascade, the chorus mean declines to silence, fluctuating between 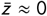 and 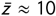 as the population devolves into chaos, ultimately resulting in total collapse (Fig. S6).

Decreasing the sensitivity of the signaling advantage to changes in population size (lower auditory sensitivity *α′*) means that signal receivers are less able to distinguish among signalers against the backdrop of a signaling population. This lower auditory sensitivity means that the fitness advantages that can be realized by signalers when populations are low are lessened. Compellingly, we observe that an increased auditory sensitivity results in period-doubling cascades and the onset of chaos occurring at much higher values of *r*_max_ (Fig. 4). As such, increased auditory sensitivity increases the stability regime of signaling populations, whether they ultimately evolve towards silence or conspicuousness. When signal receivers are unable to discern among signalers even at lower population sizes (low *α′*), silence is nearly always the end-state of selection (except when both *r*_min_ and *r*_max_ are very low), a dynamic that emerges from a higher individual fitness peak at silence (*z* = 0; Fig. 1a). It is this same dynamic that promotes an earlier onset of the period-doubling cascade leading to chaos as *r*_max_ increases (Fig. 4). That increasing auditory sensitivity promotes population stability by delaying the onset of cycles and chaos suggests that the evolution of increasingly sensitive auditory machinery may not only carry with it a reproductive advantage but promote stability of the population as a

**FIGURE 4.**
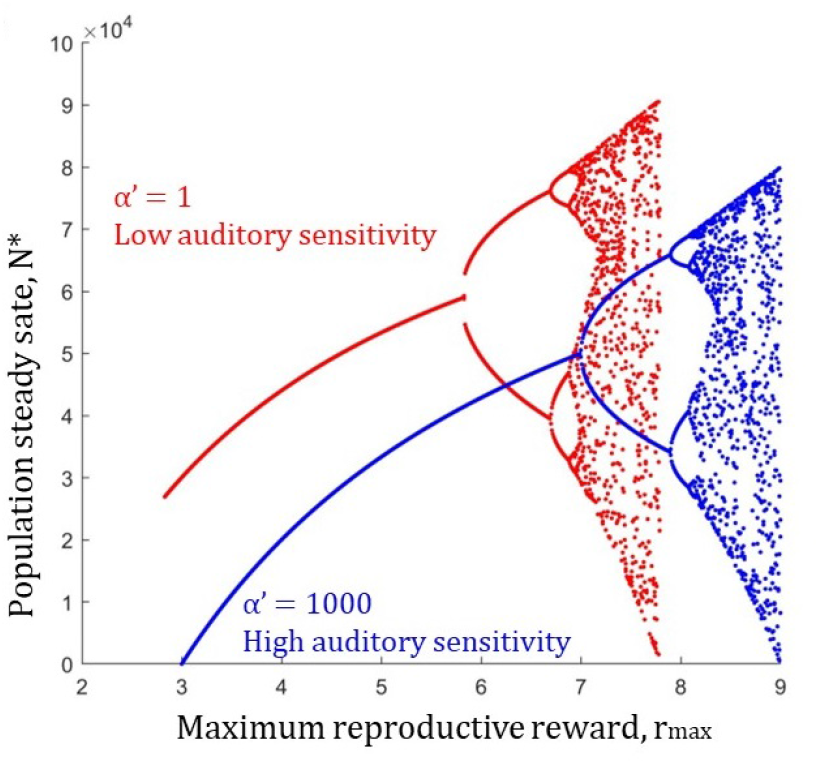
Population steady states *N*\* as a function of increasing reproductive rewards associated with signaling *r*_max_. Population steady states are shown with the auditory sensitivity of signal receivers set to low (red) and high (blue) values, resulting in the emergence of period-doubling cascades at different values of *r*_max_. Increasing the auditory sensitivity of signal receivers (higher values of *α*′) promotes the stability of signaling populations by increasing the value of *r*_max_ at which the period-doubling cascade towards chaos emerges.

That the evolution of silence versus signaling involves an interaction between the rewards of signaling and population size is supported by observations in natural systems. For example, among field crickets (*Gryllus campestris*) signaling males dominate when populations are at low densities, whereas silence emerges as a dominant strategy at high population densities, where mates are sought by alternative means (Hissmann, 1990). While this is likely a product of behavioral plasticity (a feature not present in our current approach), it demonstrates how selective feedbacks may direct evolution to alternative outcomes when behaviors are less plastic. To our knowledge, there is very limited empirical work examining the effect of changes in population density on acoustic signal evolution, and our results suggest this may be an interesting avenue for future research effort.

It is well known that changes in population densities impact the fitness of conspicuous signalers, and by extension acoustic signal evolution (Cade and Cade, 1992; Schmidt et al., 2014), however the mechanistic links between signalers and signal receivers as a function of population size is not well understood. While reproductive incentives to signal are impacted by fluctuations in the biotic and abiotic environment (Bertram et al., 2013; Hedrick and Dill, 1993; Alexander, 1975; Rowell and Cade, 1993; Boake, 1983; Burk, 1983; Cade, 1979), our results suggest that species with greater investments in the physiological machinery required to distinguish and target individual signalers may have populations less prone to large fluctuations in population size. Moreover, our results support the notion that increased investment in auditory reception may promote stability by minimizing the range of growth rates at which population cycles emerge. Of potential importance is the notion that signal receivers with lower auditory sensitivity may be more susceptible to anthropogenic noise (Orci et al., 2016; Gurule-Small, 2018). While the relationship between auditory reception complexity and the stability of acoustic signaling populations has not been explored, the potential fragility of species with naturally low auditory sensitivity - or declining sensitivity due to interference by anthropogenic noise - may have particular implications for conservation efforts.

### 2.2 The costs of signaling

The energy devoted to behaviors such as signaling is necessarily limited by that required to survive in a given environment. In less productive environments, organisms generally spend more time and energy foraging, at the cost of somatic growth and avoiding predation. In our framework, we capture the energetic costs associated with nonsignaling behaviors by *δ*_min_, the per-capita mortality of silent individuals. In contrast, we capture the energetic costs associated with signaling with *δ*_max_, the maximum per-capita mortality of conspicuous individuals.

Our results show that as the reproductive incentive Δ*r* and the cost of signaling *δ*_max_ increase together, selection favors the evolution of signaling as long as Δ*r* is sufficiently greater than *δ*_max_ (Fig. 5, *yellow region*). If the reproductive incentive is much greater than *δ*_max_, population growth quickly fuels instabilities (Fig. 5, *region d*), causing the emergence of cycles, chaos, and ultimately collapse. If the cost is much greater than the reproductive incentive, signaling is more costly than the reproductive reward, and silence emerges as the evolutionary outcome (Fig. 5, *region b*).

**FIGURE 5.**
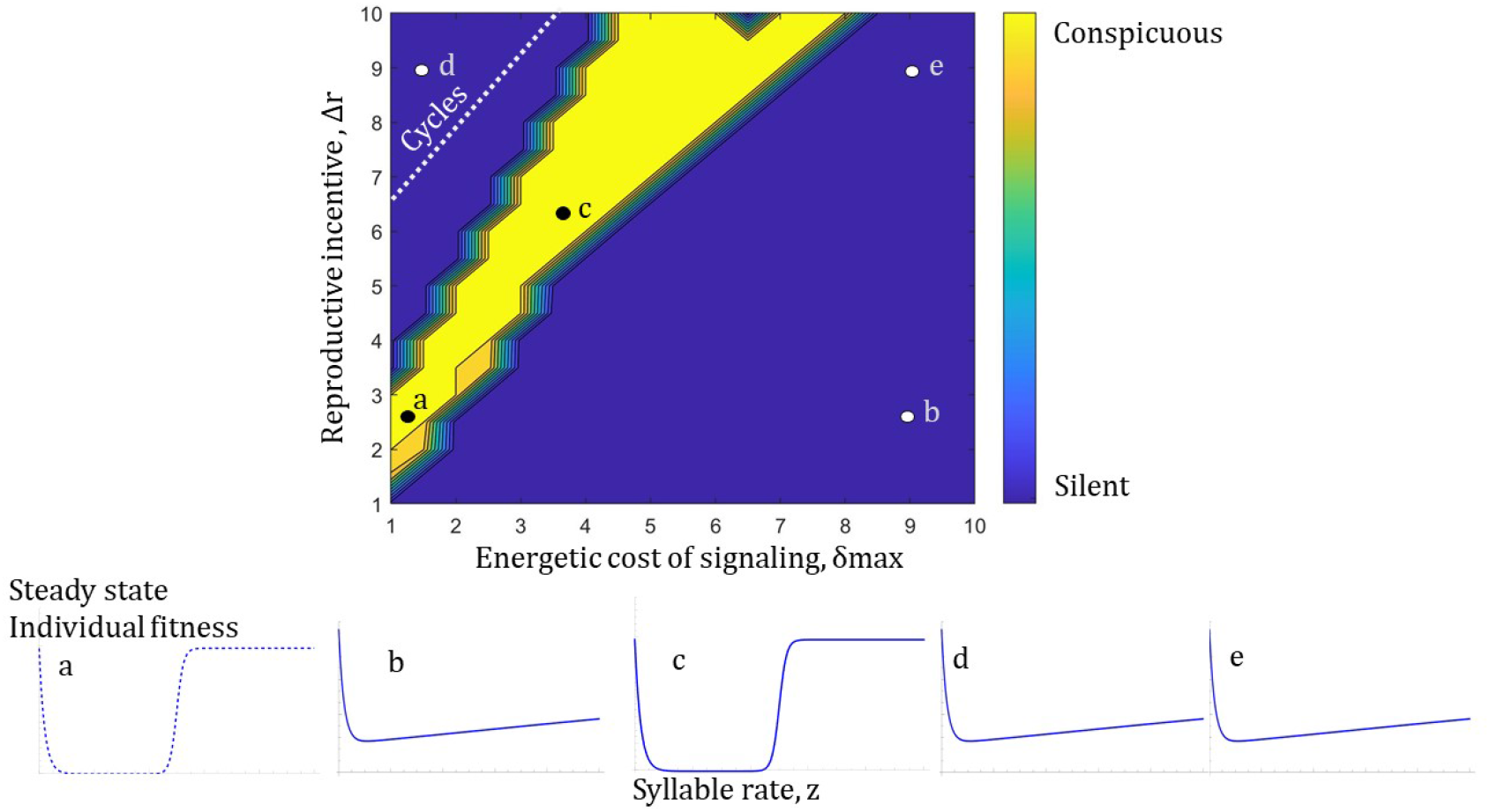
Energetic cost and reproductive incentive dependent evolution of the chorus mean steady state, 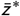. Panels (a-e) correspond to the individual fitness *w*(*z*) as a function of the acoustic trait *z* at steady state. The dashed line represents individual fitness when the corresponding population density is low.

Along with the reproductive incentive to signal, the energetic costs associated with a given environment should play a central role in shaping signaling evolution. However, while a productive environment promotes easy access to resources and enables increased allocation towards reproductive expenditures such as signaling, it alone is not sufficient to ensure that signaling is the evolutionary outcome (Tolle and Wagner Jr, 2011; Wagner Jr and Hoback, 1999; Bertram, 2002; Reifer et al., 2018; Harrison et al., 2014). Alongside lower energetic costs (lower *δ*_max_), is also the requirement that Δ*r* > *δ*_max_ to enable the evolution of conspicuous signaling. And while this requirement sets a lower bound on such an evolutionary outcome, if the reproductive rewards associated with signaling are too great, the onset of fluctuations destabilizes the population, leading to weak selection gradients and the evolution of silence, and then the more fundamental silence associated with collapse. In addition to the energetic costs of a particular environment, factors such as age, mating status, attractiveness, body size, condition, and parasite load also influence the energetic cost of signaling (Dougherty, 2021). For example, populations with greater frequencies of silent phenotypes have been observed among field cricket populations (Zuk et al., 2006), alongside the development of alternative strategies like increased movement and satellite males (Zuk et al., 2006; Rick and D Ill, 1993; Hack, 1998). The evolution of silence among species exposed to predators or parasites that cue onto acoustic signals has been examined in some detail, and where these pressures are extreme, entire populations may lose the ability to signal (Zuk et al., 2006), though a theoretical framework for this dynamic is currently lacking.

## 3 CONCLUSION

We have shown that the evolution of both conspicuous signaling and silence is driven by the trade-off between reproductive rewards, energetic costs, and the phonotactic selectivity constrained by the auditory sensitivity of signal receivers. Because the reproductive success of conspicuous signalers is greater when population densities are low (Hissmann, 1990), feedback between the size of the population and the strength of selection plays a significant role in determining which evolutionary outcome is realized. In nature, fast-growing populations can be more prone to cyclic oscillations, and perhaps chaos and extinction (Hastings et al., 2018; Melbourne and Hastings, 2008; Vortkamp et al., 2020). The likelihood of these dynamic transitions can be increased by changes in predator interactions, Allee effects, and/or mating success (Schreiber, 2003; Hastings, 2004). Moreover, the effects of predation and especially parasitism are expected to not only increase the reproductive costs of signaling, but to have compensatory effects on population size. The results of our framework suggest this may have a large influence on the evolution of signaling. Because the sensitivity of phonotactic selectivity to changes in population size largely determines whether signaling is feasible or not, the introduction of acoustic pollution in disturbed habitats may be expected to influence under what conditions signaling maximizes fitness (Bowen et al., 2020; Caorsi et al., 2017; Ortega, 2012; Laiolo, 2010). In order to gain insight into the evolution of acoustic signaling, we must understand the mechanistic connections that link signaler to signal receiver and the fitness consequences associated with individuals who strive to rise above the noise.

## Acknowledgements

We thank Emily Jane McTavish, Arnold D. Kim, Miriam Barlow, Irina Birskis-Barros, Taran Rallings, Ritwika VPS, Kinsey Brock, Laura Van Vranken, Marie-Claire Monier Chelini and Jean Phillippe Gibert for their helpful comments and suggestions. This manuscript benefited from the following UC Merced internal fellowships: Graduate Student Opportunity Program, Southern California Edison Fellowship, Dr. Donald and Effie Godbold Fellowship, QSB Summer Research Fellowship, Fred and Mitzie Ruiz Fellowship, and SNS Dean’s Fellowship.

## S4 SUPPLEMENTARY FIGURES

**FIGURE S6.**
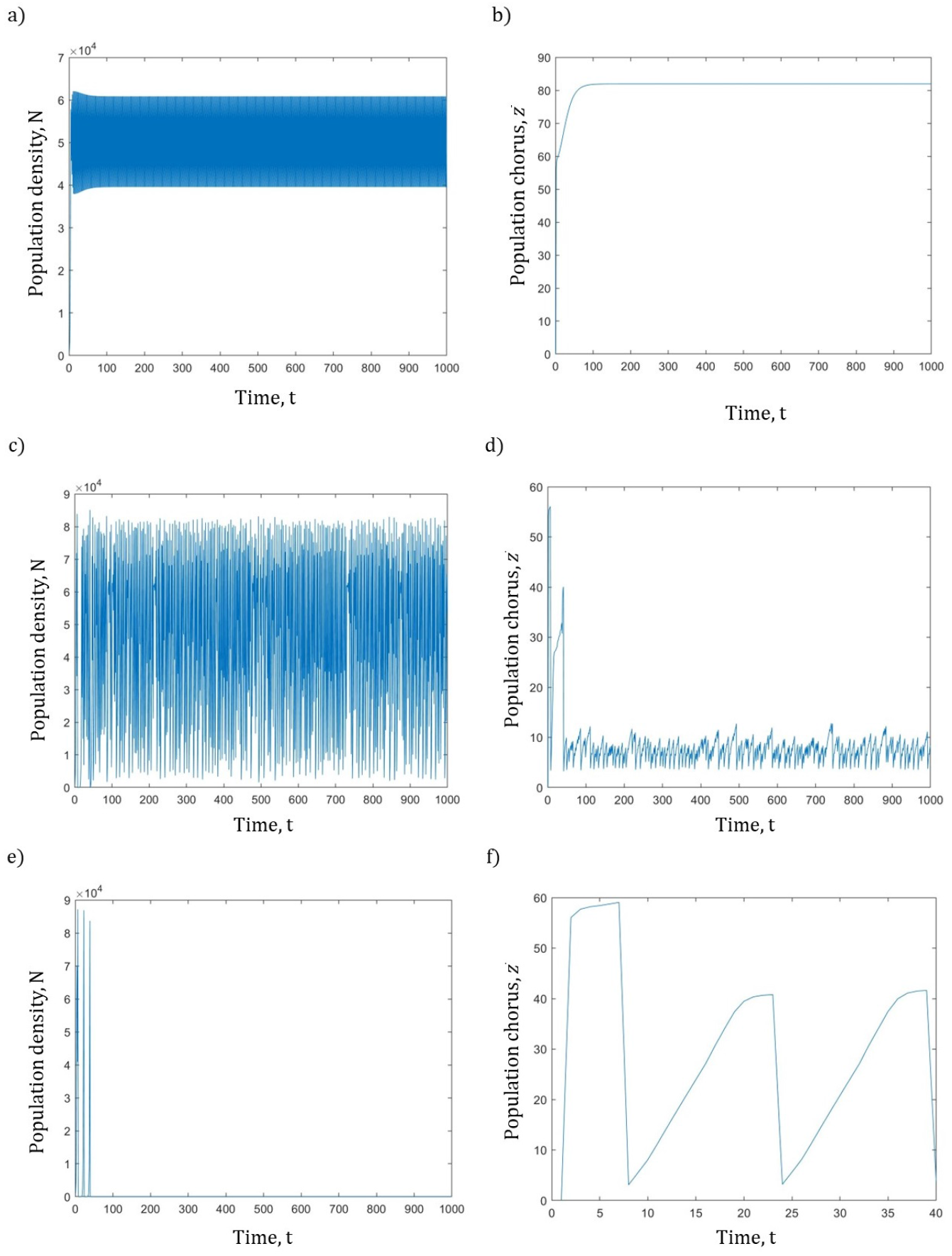
Population density and population chorus mean change over time. Left panels: Population density over time *N*. Right panels: Trait chorus mean over time 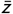 (*α*′ = 200, *r*_min_ = 1, *d*_max_ = 1, *d*_min_ = 0.01, *K* = 10^5^, *β* = 0.5, *N*_0_ = 1000, *σ_g_* = 1, and *t_max_* = 1000). a) Population cycles with high amplitude result in a conspicuous population *r*_max_ = 7.5. b) Population cycles with high amplitude result in a silent population *r_max_* = 9. c) A very high reproductive rate leads to population collapse *r_max_* = 10.

**FIGURE S7.**
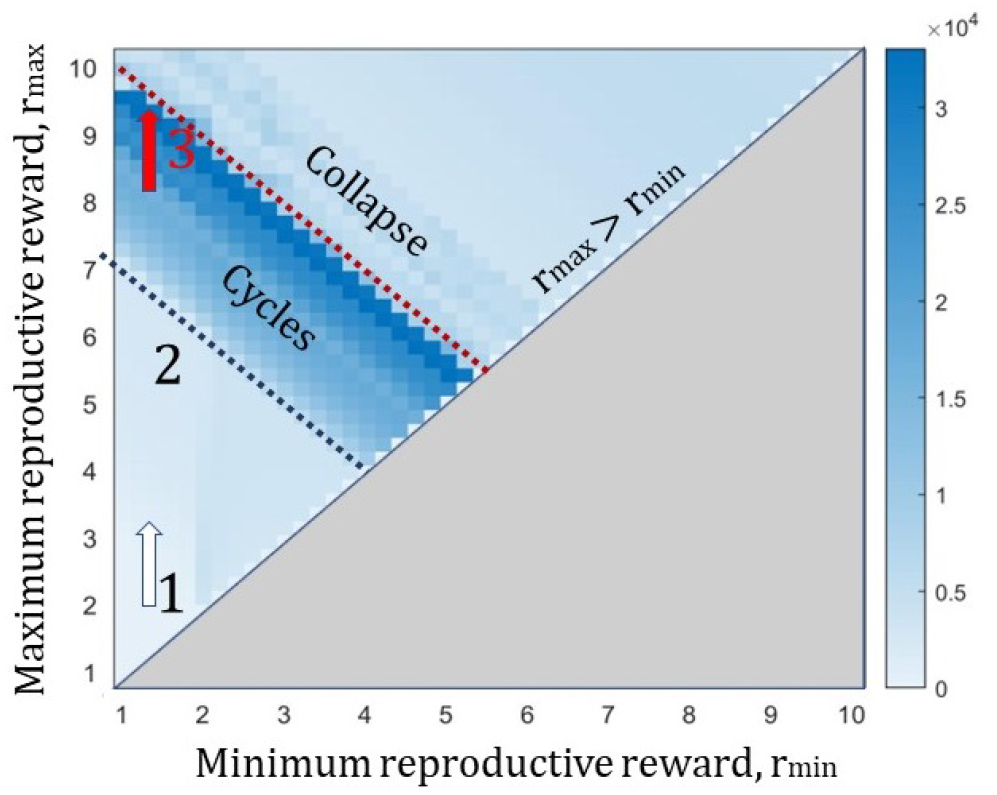
Population density steady state standard deviation for various reproductive incentives.

